# No effect of continuous transcutaneous auricular vagus nerve stimulation on the P3, the P600, or physiological markers of norepinephrine activity in an oddball and sentence comprehension task

**DOI:** 10.1101/2025.09.05.674460

**Authors:** Friederike Contier, Isabell Wartenburger, Mathias Weymar, Milena Rabovsky

## Abstract

The ERP components P3 and P600 have been proposed to reflect phasic activity of the locus coeruleus norepinephrine (LC/NE) system in response to deviant and task-relevant stimuli across cognitive domains. Yet causal evidence for this link remains limited. Here, we used continuous transcutaneous auricular vagus nerve stimulation (taVNS), a non-invasive method proposed to modulate LC/NE activity, to test whether these components are indeed sensitive to NE manipulation. Forty participants completed both an active visual oddball task and a sentence processing task including both syntactic and semantic violations, while receiving continuous taVNS at the cymba conchae in one session and sham stimulation at the earlobe in another session. We observed robust P3 and P600 effects. Crucially though, taVNS had no effect on P3 or P600 amplitude. The physiological NE markers salivary alpha amylase level and baseline pupil size were also unaffected by the stimulation, suggesting that the taVNS protocol and/or task may not have been sufficient to successfully engage the LC/NE system. Beyond the stimulation, however, exploratory analyses revealed correlations between the syntactic P600 and both the P3 and salivary alpha amylase levels, supporting the idea that the P600 might be related to both the P3 and NE. Overall, our findings do not allow for theoretical implications concerning a potential causal link between the two components and NE but highlight the need for more standardized taVNS protocols.

## Introduction

Components of the event-related potential (ERPs) have been widely studied as neural correlates of specific cognitive processes. They have traditionally been examined within domain-specific contexts – such as emotion regulation, attention in the visual oddball paradigm, or structural processing in language tasks – leading to a fragmented understanding of their functional significance. To move beyond functional description of single ERPs and toward a mechanistic understanding of these components that may cross the boundaries of cognitive subdomains, it is crucial to consider not only their cortical generators but also the neurochemical systems that support them. ERPs are assumed to reflect postsynaptic potentials from synchronized neural populations (Luck, 2014; Mitzdorf, 1994) and are modulated by neurotransmitter activity that governs neuronal excitability and synaptic transmission (Frodl-Bauch et al., 1999). However, the specific roles that different neuromodulatory systems play in shaping ERP components remain largely unexplored with some initial exceptions. For example, the locus coeruleus norepinephrine (LC/NE) system has been proposed as a key neuromodulatory mechanism underlying a family of late positive ERP components, including the P3 and P600. Here we test this hypothesis using transcutaneous auricular vagus nerve stimulation (taVNS), a noninvasive stimulation method suggested to boost the release of NE from the LC (Badran et al., 2018; Farmer et al., 2021; Frangos et al., 2015; Schneider et al., 2025). This approach allows us to probe the neurochemical underpinnings and potential overlap of these ERP components, possibly suggesting a shared, domain-general mechanism.

The P3 is a robust, centro-parietal positive ERP peaking around 300-600 ms post-stimulus, typically elicited by rare, task-relevant “oddballs” among frequent standard stimuli across domains (Donchin, 1981; Polich, 2007). It has been linked to stimulus evaluation, salience detection, attentional resource allocation, and context updating (e.g., Donchin & Coles, 1988; Kok, 2001; Nieuwenhuis et al., 2005; Verleger, 2020), but its precise functional significance remains under debate. Further, although several cortical and subcortical neural generators have been proposed – mostly focusing on parietal and frontal cortices (e.g., Bledowski et al., 2004; Halgren et al., 1998; Linden, 2005) – the neurochemical mechanisms underlying the P3 are not fully understood.

The P600, a similarly late, centro-parietal positivity peaking around 500-900 ms in response to structural anomalies in language, has long been considered a hallmark of syntactic reanalysis and repair processes during sentence comprehension (e.g., Friederici et al., 1996; Gouvea et al., 2010; Hagoort et al., 1993; Osterhout & Mobley, 1995). For example, it is observed in response to number agreement mismatches between the subject and verb of a sentence, as in “The spoilt child *<u>throw</u>*the toys on the ground.” (transl. from Dutch, Hagoort et al., 1993). However, it is also elicited by a broad range of non-syntactic linguistic anomalies, including semantic, pragmatic, orthographic, and discourse-level violations (e.g., Hoeks et al., 2004; Kuperberg et al., 2020; Regel et al., 2014) prompting proposals that the P600 reflects a more general mechanism of conflict detection or reanalysis (e.g., van de Meerendonk et al., 2009). Importantly, the P600 is even elicited by nonlinguistic violations in domains such as music, arithmetic, and visual event comprehension (e.g., Cohn & Kutas, 2015; Patel et al., 1998), challenging the assumption that the P600 is language-specific at all.

Because both the P3 and P600 are late, parietal positivities sensitive to stimulus salience and task relevance, it has been proposed that they may share common neurocognitive underpinnings (the “P600-as-P3” hypothesis, Coulson et al., 1998a, 1998b; Leckey & Federmeier, 2019; Sassenhagen & Bornkessel-Schlesewsky, 2015; Sassenhagen & Fiebach, 2019). At the same time, empirical evidence remains mixed (e.g., Osterhout et al., 1996). For instance, P600 effects can occur in the absence of explicit decision-making (Vissers et al., 2007). Thus, whether the P600 represents a domain-general late variant of the P3 or a distinct, language-specific process remains an important and ongoing debate in the field. The current study addresses this debate directly by testing for shared modulation of the two components via taVNS.

One prominent theory proposes that the P3 and P600 share a neural generator: NE release from the LC in the brain stem which plays a central role in modulating attention, arousal, and behavioral adaptation (Aston-Jones & Cohen, 2005). Such late positivities, including the P3, the late positive potential, and the P600, are assumed to reflect such phasic NE response to salient and motivationally significant events across domains (Bornkessel-Schlesewsky & Schlesewsky, 2019; Hajcak & Foti, 2020; Murphy et al., 2011; Nieuwenhuis et al., 2005; Sassenhagen et al., 2014; Sassenhagen & Bornkessel-Schlesewsky, 2015; Vazey et al., 2018). The LC-NE projects to relevant target cortices, where it promotes focused attention to relevant stimuli (neural gain, Nieuwenhuis et al., 2005) or acts as a neural interrupt signal (Bouret & Sara, 2005; Dayan & Yu, 2006).

There are indeed several functional parallels between the positivities and the LC/NE system: First, NE responses are related to motivational significance and salience as well as sensitive to stimulus probability and task relevance (Nieuwenhuis et al., 2005, 2011). Second, the LC/NE system is strongly oddball sensitive (e.g., Aston-Jones et al., 1997). Third, just like LC/NE response itself, the latency of both the P3 and P600 have been shown to align more closely with reaction times than with stimulus onsets (Bouret & Sara, 2004; Kutas et al., 1977; Sassenhagen et al., 2014; Sassenhagen & Bornkessel-Schlesewsky, 2015). This temporal alignment supports the notion that both components index similar cognitive control or salience-driven processes, potentially mediated by phasic NE. In addition, all three – the LC/NE system, the P3, and the P600 – have been implicated in long-term memory facilitation. By activating attentional networks and involvement of limbic structures, phasic LC/NE activation has been shown to promote explicit memory formation, especially under conditions of heightened salience or arousal (e.g., Aston-Jones & Cohen, 2005; Gibbs et al., 2010; Izumi & Zorumski, 1999; Mather et al., 2016; Sara, 2009). Analogously, both P3 and P600 amplitudes have been found to predict subsequent memory performance, suggesting that these components may reflect neurocognitive processes that prioritize information for encoding (e.g., Contier et al., 2025; Dunn et al., 1998; Fabiani et al., 1986, 1990; Karis et al., 1984; Neville et al., 1986; Paller et al., 1988).

There is also ongoing research attempting to directly link the positivities to activity of the LC and its correlates. For example, unit recordings suggest simultaneous firing of LC neurons and P3 during oddball tasks in macaques (Swick et al., 1994). As measuring the LC directly remains challenging in humans, much research focuses on the task-evoked pupillary response, a common non-invasive marker of NE (e.g., Joshi et al., 2016; Liu et al., 2017; Murphy et al., 2014; Rajkowski et al., 1993; Sterpenich et al., 2006). While earlier studies failed to find covariation between the pupil and P3 (Hong et al., 2014; Kamp & Donchin, 2015; LoTemplio et al., 2020; Murphy et al., 2011), recent research suggests that it can be uncovered if response-related activity is taken into account (Chang et al., 2024; Menicucci et al., 2024). Contier et al. (2024) also found first evidence that the amplitude of the P600 covaries with the task-evoked pupillary response, particularly its temporal derivative. Linking the late, positive ERP components and pupil dilation remains challenging because the two operate on different timescales and the pupil reflects multiple overlapping processes only some of which may be related to the late positive ERPs. Additionally, late positive ERP components and the pupil dilation are similarly sensitive to task conditions (e.g., oddball vs. standard trials), which can be seen as a confound (see Contier et al., 2024, for discussion), making it difficult to isolate a shared underlying mechanism such as LC/NE activity.

Beyond correlation, it is crucial to establish a causal relationship between the two positivities and the LC/NE system. Previous research suggests that the P3 can indeed be modulated by interventions that alter NE release, such as pharmacological manipulations (e.g., Joseph & Sitaram, 1989; Pineda & Westerfield, 1993). Together with lesion studies in rodents and monkeys (e.g., Ehlers & Chaplin, 1992; Pineda et al., 1989), these studies indeed suggest a causal link between the P3 and NE, while such intervention research regarding the P600 is lacking thus far.

TaVNS has emerged as a promising non-invasive technique to engage central neuromodulatory pathways, with a particular emphasis on the LC/NE system (Badran et al., 2018; Farmer et al., 2021; Frangos et al., 2015; Szeska et al., 2025). The method involves electrical stimulation of the auricular branch of the vagus nerve, typically at the cymba conchae or tragus, which are known to be innervated by afferent vagal fibers (Badran et al., 2018; Peuker & Filler, 2002). These fibers project to the nucleus tractus solitarius in the brainstem, which relays signals to key neuromodulatory centers, such as the LC (e.g., Frangos et al., 2015; Kawai, 2018). Converging evidence from animal models and neuroimaging supports the notion that (ta)VNS can engage the LC and modulate central NE activity (e.g., Dorr & Debonnel, 2006; Ludwig et al., 2023; Schneider et al., 2025; Yakunina et al., 2017). In humans, taVNS has been found to affect task-evoked pupil responses (e.g., D’Agostini et al. 2023; Lloyd et al., 2023; Ludwig et al., 2023; Pervaz et al., 2025; Sharon et al., 2021; Skora et al., 2024; Villani et al., 2022; but see e.g., Keute et al., 2019; C. M. Warren et al., 2019; Burger et al., 2020), salivary alpha-amylase levels (e.g., Giraudier et al., 2022; Schneider et al., 2025; Ventura-Bort et al., 2018; C. M. Warren et al., 2019), heart rate variability (Kaduk et al., 2025) as well as electrophysiological processing in attention and memory tasks (e.g., Collins et al., 2021; Giraudier et al., 2020; Ventura-Bort et al., 2021; Ventura-Bort et al., 2025), all of which are influenced by LC/NE activity. It is important to note that taVNS may not act exclusively through the LC/NE system. In addition to LC/NE-related mechanisms, taVNS has been shown to modulate GABAergic inhibitory processes (Boscarol et al., 2025) and to influence earlier stages of attentional and cognitive processing, as reflected in modulations of components such as the P2 (Jelincic et al., 2025) which have also been interpreted within LC/NE-based attentional frameworks. Thus, while LC/NE mechanisms are central to many current taVNS models, other neuromodulatory pathways may also contribute to its effects.

Several studies have tested whether taVNS can modulate the amplitude of the P3 but findings to date have been mixed. Some studies have reported increased P3 amplitudes under taVNS, particularly in auditory or visual oddball paradigms, although sometimes restricted to specific types of oddball stimuli (Giraudier et al., 2024; Gurtubay et al., 2022; Rufener et al., 2018; Ventura-Bort et al., 2018; C. V. Warren et al., 2020). For instance, Ventura-Bort et al. (2018) found that while there was no overall effect of taVNS on the P3, modulations were observed, particularly for easy targets (c.f., Giraudier et al., 2024). Others have failed to observe any reliable effects (D’Agostini et al., 2022; Fischer et al., 2018; Gadeyne et al., 2022; C. M. Warren et al., 2019). Thus, the literature remains inconclusive, motivating further replication, especially when combined with physiological validation measures. Importantly, evidence of potential taVNS effects on the P600 are completely lacking. If the P600 also reflects a phasic LC/NE response to salient stimuli, then increasing LC/NE activity via taVNS should amplify or modulate the P600, similar to how it might affect the classic oddball P3.

Here we thus investigated whether the P3 and P600 are causally linked to the LC/NE system using taVNS. While receiving continuous taVNS (one session) and sham stimulation (another session), participants completed an active oddball paradigm aimed to elicit the P3 as well as a sentence processing task including both syntactic and semantic violations aimed to elicit the P600. Since taVNS is assumed to innervate the LC/NE system, the amplitude of both the P3 and P600 should be enhanced under taVNS compared to sham if they reflect LC/NE activity. In addition to the P3 and P600, we also examined the N400 component as a theoretically informative comparison. The N400 and P600 are often considered functionally and neurophysiologically distinct in language processing (e.g., Kuperberg et al., 2020; Rabovsky & McClelland, 2020): the N400 is typically linked to semantic prediction and integration, while the P600 is associated with more controlled, attention-sensitive processes.

Because only the latter has been hypothesized to depend on phasic LC/NE activity, we did not expect taVNS to modulate the N400. Including both components allowed us to assess the specificity of potential taVNS effects on late positivities in language processing. To confirm that taVNS (but not sham) indeed affected the LC/NE system, we assessed salivary alpha amylase levels, a putative autonomic NE correlate (e.g., Bosch et al., 2011; Nater & Rohleder, 2009) that has previously been shown to be modulated by taVNS (e.g., Giraudier et al., 2022; Schneider et al., 2025; Ventura-Bort et al., 2018; C. M. Warren et al., 2019). To contribute further to evidence for potential taVNS markers, we additionally measured baseline pupil size at several points in the study, as this NE-associated marker (e.g., Joshi et al., 2016; Laeng et al., 2012; Reimer et al., 2016) has to the best of our knowledge not been successfully manipulated via taVNS (e.g., Burger et al., 2020a, 2020b; D’Agostini et al., 2022; Keute et al., 2019; Skora et al., 2024).

## Method

The study including methods, hypotheses, and statistical models was pre-registered on the Open Science Framework (https://osf.io/fgbxs/?view_only=743d5d402e5249de8d4f2ed753dd5bb4). Any deviations from this plan are explicitly stated. The linked OSF project (https://osf.io/qfhab/?view_only=510598a63fe542fe8e4f6758585841b9) also hosts all pre-processing and analyses scripts as well as the pre-processed data.

### Participants

Forty naïve volunteers participated in the experiment (35 female, 5 male). Three participants had to be excluded and replaced since they did not complete both sessions (preregistered data exclusion criteria). Mean age was 24.1 years (*SD* = 5.5, range = 18-38). All participants were native speakers of German and right-handed according to the Edinburgh Handedness Inventory (Oldfield, 1971; 12-item version; mean laterality quotient: 86, range: 41-100). Participation was compensated monetarily or with course credit. Each individual provided written informed consent for a protocol approved by the ethics committee of the host institution (No. 85/2020 amendment). Participants were pre-screened online for the following exclusion criteria: Impaired and uncorrected vision, nystagmus, pregnancy, metal close to the left ear, chronic or acute medication, neurological or mental disorders, chronic illnesses (endocrine, metabolic, cardio-vascular, pulmonary, renal), metal implants, pumps, or pacemakers, history of migraine, epilepsy or brain surgery, and/or speech/language impairments.

### Procedure

The study consisted of two experimental sessions, which took place seven days apart, at the same time of the day. In one session, participants received the vagus stimulation and in the other session the sham stimulation (order counterbalanced across participants). Participants were not aware of the manipulation (single-blind study). Apart from the stimulation condition, the two sessions were identical (Figure 1A below). Testing took place in a sound-proof and electrically shielded booth with dimmed light. After study information, consent, and preparation of the EEG cap, the NE markers salivary alpha amylase levels and baseline pupil size were measured (“pre”). Subsequently, the stimulation electrode was applied, intensity adjusted, and device turned on (see below). Then, the two tasks (oddball task, sentence processing task) proceeded, in counterbalanced order (but kept constant across sessions for each participant). Between the tasks, there was a short break in which salivary alpha amylase levels and baseline pupil size were measured again (“post1”). As soon as the second task was finished, the stimulation device was turned off and salivary alpha amylase levels and baseline pupil size were measured one last time (“post2”). Timing of both tasks (both stimulus presentation times and breaks) were controlled as much as possible in order to keep the duration of the experimental session under stimulation as constant as possible. Each experimental session lasted approximately 40 minutes. At the end of each session, participants filled out a questionnaire about potential adverse effects of the vagus nerve stimulation: On a 7-point scale ranging from 1 (not at all) to 7 (very much), they rated how much they felt the following sensations during the stimulation: headache, nausea, dizziness, neck pain, neck muscle contractions, stinging sensation, skin irritation on the ear, fluctuations of concentration, fluctuations of feelings, or other unpleasant feelings (Giraudier et al., 2020, 2025; Ventura-Bort et al., 2018). At debriefing after the second session, participants learned about the broader theoretical goal of the study as well as that it involved two stimulation settings for comparison more generally. None of the participants indicated that they knew or guessed the true nature of sham vs. active stimulation during the experiment.

**Figure 1.**
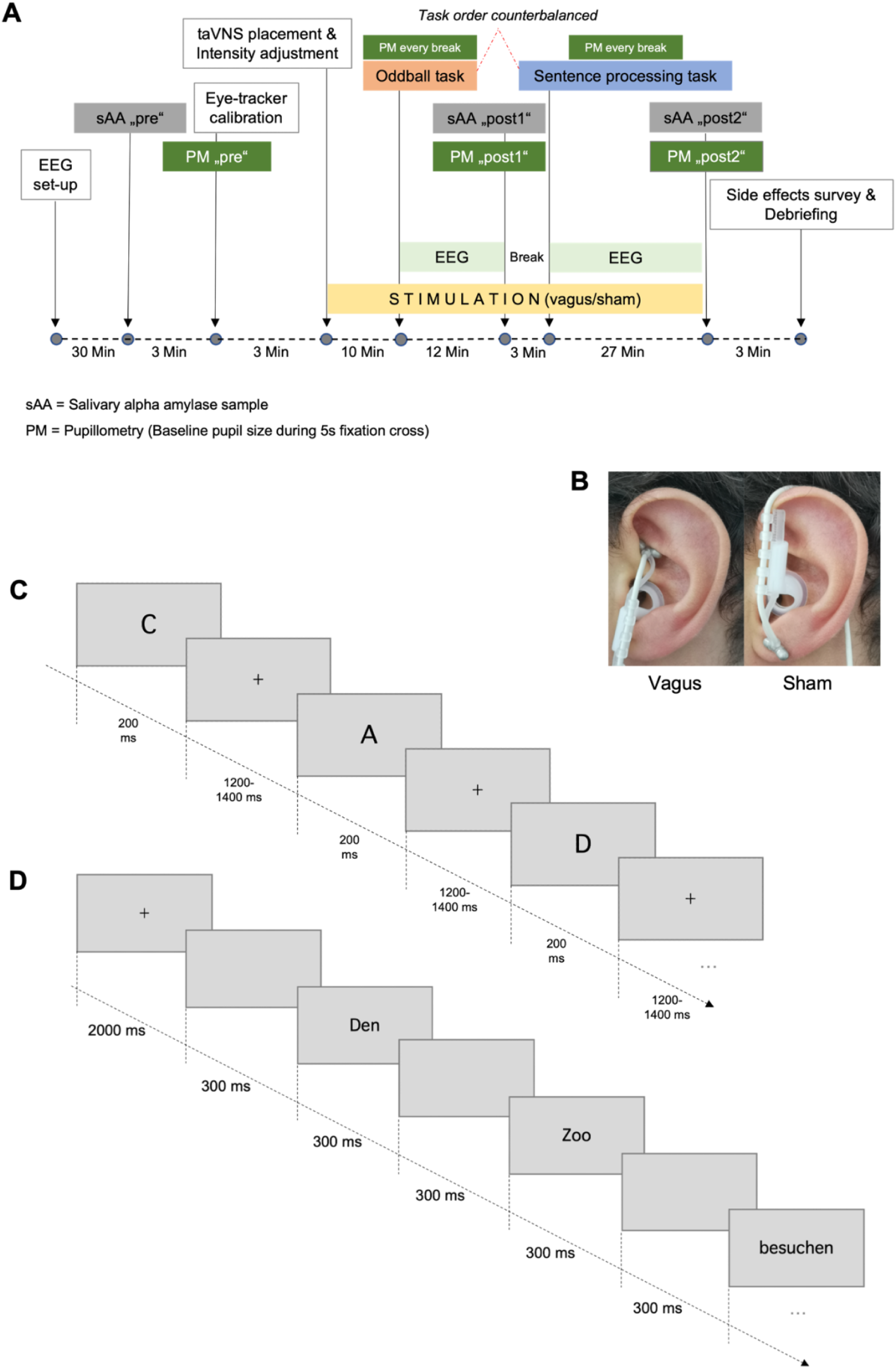
(**A**) Experimental procedure, identical across the two sessions and stimulation conditions. Stimulation was either vagus (one session) or sham (another session). **(B**) Set-up of taVNS stimulation device in the vagus (left) and sham (right) condition (from Giraudier et al., 2020). (**C**) Stimulus presentation in the oddball task, with an example stimulus sequence in the block in which “D” was the target letter. (**D**) Word-by-word stimulus presentation of one trial in the sentence processing task.

### Transcutaneous auricular vagus nerve stimulation (taVNS)

The taVNS stimulation device consisted of two titan electrodes attached to a stabilizing mount and wired to a stimulation unit (CMO2, Cerbomed, Erlangen, Germany). The mount was always placed in the left auricle, but the location of the electrodes differed between conditions (Fig. 1B). In the vagus condition, the electrodes were positioned at the cymba conchae, which is exclusively innervated by the vagus nerve (Peuker & Filler, 2002). In the sham condition, the electrodes were rotated so that they touched the center of the earlobe (e.g., Frangos et al., 2015; Ventura-Bort et al., 2018). Thus, in the sham condition, participants had the same tingling sensation as in the vagus condition, but the stimulation did not affect the vagus nerve, since the earlobe is assumed not to be innervated by the vagus nerve (Peuker & Filler, 2002). Conservatively, stimulation was applied on the left ear to avoid potential cardiac side effects, although recent research does not suggest any risks even when using the right ear (e.g., Gentile et al., 2025; Tan et al., 2025). The device emitted continuous stimulation at a pulse width of 200-300 µs at 25 Hz for the duration of both tasks (appr. 40 minutes). Since the stimulation sensation varies across individuals, we adjusted the intensity of the stimulation to ensure that all participants had the same sensation during the stimulation, above the detection, but below the pain threshold (c.f., Ventura-Bort et al., 2018). In an adjustment procedure using a scale from 1 (no sensation) to 10 (pain), we determined the intensity with which the participants judged the sensation as an 8 (maximum tingling, without discomfort). The mean intensity in the vagus condition was 0.99 mA (*SD* = 0.541, range = 0.4-2.4) and in the sham condition 1.088 mA (*SD* = 0.649, range = 0.4-3.1). Statistically, intensities did not significantly differ between stimulation conditions (*t*(39) = −1.563, *p* = .126, d = 0.163).

### Tasks & materials

Stimuli of both tasks were presented visually on a light grey computer screen in black Geneva font using MATLAB’s Psychtoolbox-3 (Brainard, 1997), at a distance of 60 cm from the participants.

#### Oddball task

To elicit the P3, we used an active, visual oddball task specifically designed to elicit a strong ERP effect while diminishing physical confounds (Kappenman et al., 2021). The task is to identify rare target stimuli (‘oddballs’) across many frequent non-targets (‘standards’).

In each instance of the task, five letters served as stimuli, each presented with a probability of .2. There were five blocks of trials and for each block, another of the five letters was designated to be the target letter (oddball) for that block. Thus, in each block, the oddball probability was .2 and the standard probability was .8, while each letter served both as a target and non-target across the whole task. Within a block, letters were presented in random order, with the restriction that targets were never followed by targets. The order of blocks was also randomized. In each session and thus, stimulation condition, we presented a total of 200 trials (40 targets and 160 non-targets), divided into five blocks of 40 trials each.

In one session, the stimulus letters were A, B, C, D, and E and in the other, K, L, M, N, and O (counterbalanced across stimulation conditions and sessions). Stimulus letters were presented in the center of the screen (height: 6.676 degrees of visual angle) for 200 ms (Fig. 1C). During the inter-stimulus interval (ISI), a fixation cross was shown for 1200-1400 ms (jittered in steps of the screen refresh interval of 16.667 ms using a rectangular distribution).

Participants’ task was to respond as quickly and accurately as possible whether the presented letter is the target or non-target for that particular block. Two marked (control) keys served as response buttons for ‘left’ and ‘right’ responses, each pressed by the index finger of the respective hand. Stimulus-response mapping (right key target & left key non-target, or vice versa) was counterbalanced across participants, but held constant within participants across tasks and sessions (see below). Written instructions were presented at the beginning of the task and reminders after each break within the task. Before the experimental blocks, participants practiced the task using a set of letters different from the experimental letters (27 trials) and were only allowed to proceed with above-chance accuracy. Between blocks, participants had a short break to rest their eyes (20 seconds). The task lasted approximately 12 minutes.

#### Sentence processing task

This task was a classic sentence processing task in which sentences are presented word by word and participants are prompted to make an acceptability judgement after each sentence. We used German sentences by Sassenhagen & Fiebach (2019, Exp. 2), which contain 50% correct sentences, 25% syntactic violations and 25% semantic violations.

Sentences were simple, plausible subject-verb-object (S-V-O) or object-verb-subject (O-V-S) sentences. For the syntactic violation, a second version of each sentence with an agreement violation had been constructed by exchanging the verb or pronominal subject (example 1 below) for a morphosyntactic ill-fitting one from another sentence (thereby creating an agreement violation). Although this constitutes a morphosyntactic violation, we will henceforth call it “syntactic” violation for simplicity. Semantic violations had been created by either exchanging a noun (example 2 below) or verb between two experimental sentences rendering each of the sentences highly implausible.

1. „Den Zoo besuchen_[pl]_ **<u>sie</u>**_[pl]_ **<u>/ *er</u>**_[sg]_ am Montag” [The zoo visit **<u>they / *he</u>** on Monday]
2. “Der Chirurg operiert **<u>den Leistenbruch/*die Butter</u>** mit Vorsicht” [The surgeon operates the **<u>rupture/*butter</u>** carefully]

The critical target words containing the violation appeared at variable positions within the sentence (typically at word 3, 4, or 5) and included different word classes such as verbs, nouns, and pronouns. This variability ensured that participants could not predict the position or lexical type of an upcoming anomaly. For more details on the construction of these sentences, see Sassenhagen & Fiebach (2019, Exp. 2). We also added 30 correct filler sentences in the same fashion as correct control sentences to increase the proportion of correct sentences. The final set comprised 350 sentences: 320 experimental sentences (80 syntactic violations and 80 correct control counterparts; 80 semantic violations and 80 correct control counterparts) as well as 30 correct filler sentences. These sentences were divided into 2 lists, which were counterbalanced across participants. This resulted in a classic latin-square design where each participant sees only one version of each sentence, but across participants, each sentence is presented in both its correct control and violation version equally often. These lists were then further divided into two parts, one for each experimental session (order counterbalanced across sessions and stimulation conditions). Thus, in each session (and thus, stimulation condition), participants were presented with 175 sentences of which 46% contained some kind of violation (23% syntactic, 23% semantic).

Within each session, sentences were presented in random order, with the only constraint that violation sentences of a certain type were never directly followed by another violation sentence of the same type. Each trial (see Figure 1D) started with a fixation cross, shown for two seconds. Single words of the sentence were then presented consecutively, each for 300 ms with a 300 ms blank inter-word interval. If a non-target word’s length exceeded 12 letters, 16.667 ms (one screen refresh interval) were added to its presentation duration for each additional letter (for comparable approaches, see e.g., Contier et al., 2024; Hodapp & Rabovsky, 2021; Nicenboim et al., 2020). Words had a height of 1.41° degrees of visual angle. 300 ms after the offset of the final word, a question mark appeared, prompting the participant to decide as fast and accurately as possible whether the presented sentence was syntactically or semantically anomalous. Thus, participants made their acceptability judgment only after the sentence ended to avoid overlap between language-related EEG activity and motor preparation or response execution. Reaction times therefore reflect the time required to make the final acceptability decision rather than immediate processing of the violation. Two marked (control) keys on the computer keyboard served as response buttons for ‘left’ and ‘right’ responses, each pressed by the index finger of the respective hand. Stimulus-response mapping was counterbalanced across participants but held constant in both tasks and sessions within each participant. Thus, the same key, either the left or right control key, was assigned for targets in the oddball task and violations in the sentence processing task and vice versa. The question mark was shown until button press (but max. for 2500 ms). Participants received no feedback. The next trials started 300 ms after the response.

Short rest breaks were provided every 29 trials (approx. 20 seconds). Prior to the start of the experimental sentences, participants were provided with written instructions. Then, participants practiced the task with nine additional sentences including correct control, syntactic violation and semantic violation sentences and were only allowed to proceed with above-chance accuracy (with one chance of repetition). The task lasted approximately 27 minutes.

### Data acquisition & processing

#### EEG data

In both tasks and sessions, we recorded participants’ electroencephalogram (EEG) using 32 passive Ag/AgCl electrodes (actiCHamp Plus system, Brain Products GmbH, Gilching, Germany), spaced according to the international 10-20 system. Impedances were kept below 5 kOhm if possible. The ground electrode was positioned at FCz and the reference electrode at the right mastoid. Right-side VEOG as well as bilateral HEOG electrodes were positioned to capture eye movements and blinks. The EEG was recorded at a sampling rate of 1000 Hz (with a bandpass filter of .016-250 Hz and time constant of 10s).

Offline, we preprocessed the data in *MATLAB* R2020a (The MathWorks Inc., 2020), using the *EEGLAB* toolbox (Delorme & Makeig, 2004). Data were first downsampled to 500 Hz and re-referenced to the average of the left and right mastoid. We then used independent component analysis (Infomax ICA, Jung et al., 2001) to correct ocular artifacts. ICA was trained on epochs spanning −0.5 to +2s (oddball task) and −1 to +2s (sentence processing task) relative to stimulus/target word onset after artifact rejection (removal of epochs exceeding +150/-150 mV). These epochs were extracted from continuous data downsampled to 125 Hz and filtered with a 1-30 Hz Butterworth filter. We removed independent components that the *IClabel* plug-in (Pion-Tonachini et al., 2019) identified to be eye-related with a probability equal to or larger than 30%. We additionally removed (max. three) channel noise components or components clearly originating from the taVNS device (predominantly 25 Hz activity on left side) based on visual inspection.

Corrected, continuous data were high-pass filtered (0.1 Hz, two-pass Butterworth with a 12 dB/oct roll-off) and low-pass filtered (30 Hz, two-pass Butterworth with a 24 dB/oct roll-off). Residual noise caused by the taVNS device was filtered out using EEGLAB’s CleanLine plug-in (at 25Hz + harmonics). Bad channels exceeding an absolute z-score of 3 regarding voltage across channels were interpolated using a spherical spline function. Data were then epoched from −200 to 800 ms (oddball task) and from −200 to 1100 ms (sentence processing task) time-locked to stimulus/target word onset and baseline corrected relative to the 200 ms interval preceding this onset. Artifactual trials exceeding +/-75 mV were automatically removed.

As preregistered, error trials (i.e., trials in which participants responded incorrectly) were excluded. Table 1 displays the number of trials per task and condition combination after data preprocessing. In none of the stimulus conditions did the number of trials significantly differ between stimulation conditions (all *p*’s > .13). ERP amplitudes were quantified as mean voltage values extracted from predefined component-specific time windows and electrode clusters on each trial and analyzed using linear mixed-effects models. Although the models operate on trial-wise amplitude estimates, they target condition-level effects on canonical ERP components (P3/P600), analogous to traditional averaged-ERP analyses, while additionally accounting for trial-, participant-, and (where applicable) item-level variability. Trial-wise and participant-wise variability in the data entering the models is illustrated in Supplementary Figure A1. Data for both late, positive components were analyzed in a parietal region of interest (ROI: CP1, CPz, CP2, P3, Pz, P4, PO3, POz, PO4) within a 300-600 ms (P3) and 600-900 ms (P600) time window. These spatial ROIs and time windows had been chosen based on the topography and time where each of the components had been shown to be largest in the respective task in previous studies (Contier et al., 2024; Kappenman et al., 2021; Sassenhagen & Fiebach, 2019). N400 amplitudes (in response to semantic violations and controls) were computed within a 300-500 ms time window. As preregistered, we computed N400 amplitudes within two different ROIs: For comparability, the parietal region identical to the one for the P3 and P600 mentioned above and a centro-parietal channel cluster more commonly chosen for the N400 (Cz, CP1, CPz, CP2, Pz; see e.g., Hodapp & Rabovsky, 2021; Kuperberg et al., 2020).

**Table 1.**
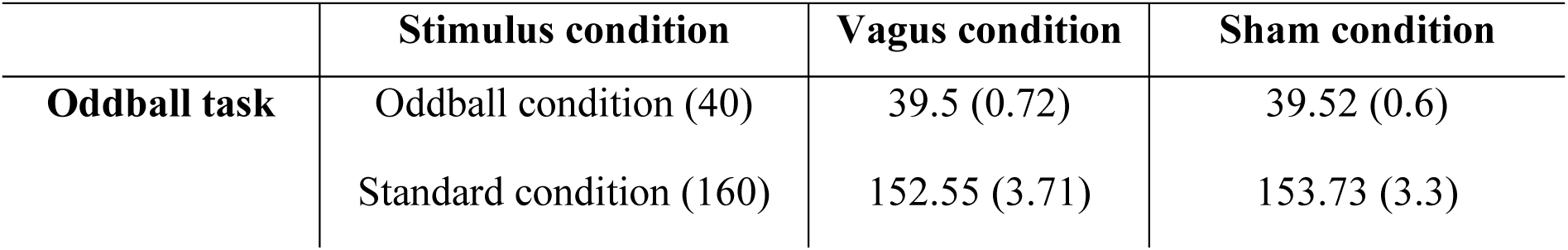
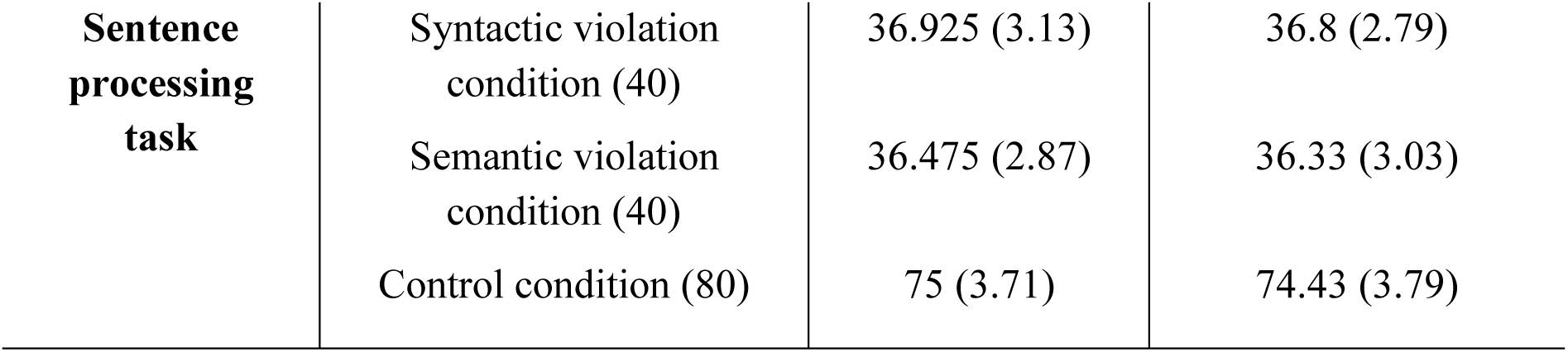
Mean (SD) number of trials per participant in each task, stimulus condition, and stimulation condition after data preprocessing. Number of presented trials in parentheses behind the stimulus condition.

|  | Stimulus condition | Vagus condition | Sham condition |
| --- | --- | --- | --- |
| Oddball task | Oddball condition (40) | 39.5 (0.72) | 39.52 (0.6) |
|  | Standard condition (160) | 152.55 (3.71) | 153.73 (3.3) |
| **Sentence processing task** | Syntactic violation condition (40) | 36.925 (3.13) | 36.8 (2.79) |
|  | Semantic violation condition (40) | 36.475 (2.87) | 36.33 (3.03) |
|  | Control condition (80) | 75 (3.71) | 74.43 (3.79) |

#### Salivary alpha amylase levels

As our main manipulation check, salivary alpha amylase (sAA) levels were collected that are proposed to correlate with NE activity (Bosch et al., 2011; Nater & Rohleder, 2009). Saliva was collected at three points during each session to investigate potential effects of the stimulation: Before the stimulation was turned on (pre), during the stimulation in the break between the two tasks (post1) and right after the stimulation was turned off (post2). Saliva was collected using the spitting method (Hodgson & Granger, 2013): Participants were instructed to pool saliva in their mouth for about one minute and then spit it into a tube using a plastic straw. Samples were stored at −18 degrees Celsius and alpha amylase levels determined via immuno-assay by *Dresden Lab Services GmbH*. Data from three participants had to be excluded since they had at least one sample in which sAA levels could not be determined. Alpha amylase levels were not normally distributed (skewness = 1.549, *SE* = 0.162) and were thus log-transformed.

sAA levels were also used to explore whether the effect of stimulation on ERP amplitudes depends on whether the stimulation actually affected the NE system in a particular participant. For these analyses, we calculated sAA “change” values that quantify how much sAA levels changed from baseline (pre). For each participant and session (i.e., stimulation condition), sAA values before stimulation onset (pre) were subtracted from values during (post1) and after (post2) the stimulation. Positive change values thus indicate an increase of sAA levels and negative values a decrease of sAA values compared to the time before the stimulation was started.

#### Baseline pupil size

As a second potential manipulation check, pupil baseline was measured since pupil size has been proposed to be influenced by the LC/NE system. Pupil size was recorded briefly before and after each task, and during short breaks, providing a measure of tonic (baseline) pupil diameter. Continuous recording during task performance was not implemented. Thus, no phasic, stimulus-evoked pupil responses were obtained (see Contier et al., 2024, for research correlating the late positive ERP components to the task-evoked pupillary response).

Pupil size was measured for 5000 ms while participants sat quietly and looked at a fixation cross. Just like saliva, we were mainly interested in pupil size at three timepoints in each session: Before the stimulation was turned on (pre), during the stimulation in the break between the two tasks (post1) and right after the stimulation was turned off (post2). In addition, we aimed to capture any potential more fine-grained or short-lived effects of the stimulation on pupil size by measuring the pupil size during every short break (i.e., every 3-4 minutes) during the tasks starting right before the stimulation was turned on and finishing right after it was turned off, resulting in 15 measurements including the three main timepoints of interest (pre, post1, post2). Pupil size of both eyes was measured using a Tobii Pro Nano mounted below the stimulus monitor at 60 Hz. The pre and post1 measurement were preceded by a 5-point calibration procedure (see e.g., Mckinnon et al., 2020; Trueswell et al., 2013).

Immediately preceding each calibration, the software prompted the participant to adjust their posture until the eye-tracker indicated a 60 cm viewing distance (within the recommended operating range of 45–85 cm). No chin rest was used, since it could compromise the combined taVNS-EEG set-up which involves many cables around the head. Although a chin rest can reduce head movement and maintain a perfectly fixed position, the Tobii Pro Nano’s design allows reliable tracking within a modest head-movement range and the short, intermittent pupil recordings (rather than continuous tracking) minimized the risk of positional drift across measurements. Offline, pupil size data were pre-processed in MATLAB using the publicly available pipeline by Kret and Sjak-Shie (2019), which performs automated artifact correction by removing dilation-speed outliers, trend-line deviations, and secluded samples. Pupil size was then averaged over the left and right pupil, low-pass filtered (cutoff at 4Hz) and interpolated. Before interpolation, epochs were excluded that had < 50% data points left. Our final measure for statistical analyses was the mean pupil size at each timepoint.

### Statistical analyses

For the primary, preregistered analyses of ERPs and the potential effect of the stimulation, we performed linear mixed-effects model (LMM) analyses using the *R* package *lme4* (R core Team, 2018). This approach accounts for subject-specific and item-specific variability and utilizes all trials, increasing statistical power (Baayen et al., 2008). These models had the following fixed effects structure:

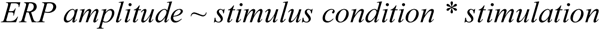

All models were built using trial-by-trial data. In these models, both categorical regressors were sum coded^1^ (Schad et al., 2018). In addition, these models included a random effects structure. In all models, we included random intercepts and slopes for the main effects and interaction between the fixed effects per subject. In models with sentence processing task data, we additionally included random intercepts and slopes for the main effects and interaction between the fixed effects per sentence. Following the recommendations by Barr et al. (2013), we tried to fit this maximal random effect structure but reduced its complexity successively until the model converged. As pre-registered, we first increased the number of iterations to 10000, then introduced an optimizer (“bobyca”), and then removed correlations between random intercepts and slopes and if necessary, entire random slopes (see analysis code on OSF for final model structures). The significance of fixed effects was determined via likelihood ratio tests to compare the fit of the model to that of a reduced model lacking the respective fixed effect but including all remaining fixed effects as well as the same random effect structure. To further investigate potential interaction effects, we conducted sub-group contrasts using the *emmeans* package (Lenth et al., 2023) with Bonferroni correction for multiple comparisons. We preregistered that if no effect of stimulation can be found, we would explore any shorter lived differences between vagus and sham stimulation on ERP amplitude. These tests compared subject-wise averaged ERP amplitudes between vagus and sham conditions in response to oddball/violations across all timepoints of the respective epoch. Clusters of consecutive timepoints exceeding a two-sided threshold (p < .05, Bonferroni-corrected) were tested against a null distribution of maximum cluster masses (5000 permutations, paired-sample t-tests).

To analyze potential effects of the stimulation on the NE markers, we conducted 2 (stimulation) x 3 (timepoint) repeated measures ANOVAs. As preregistered, we generally used the standard *p* < .05 criterion for determining if the likelihood ratio tests suggest that the results are significantly different from those expected if the null hypothesis were correct.

### Preregistered hypotheses

#### Norepinephrine markers

If taVNS affects NE activity, two assessed NE markers (salivary alpha amylase and baseline pupil size) should increase more strongly or decrease less (from pre-stimulation to post-stimulation) under vagus than sham stimulation. This effect could arise regarding either of the two post-stimulation timepoints (in between tasks and after offset of the stimulation). If no such interaction effect on pupil baseline can be detected at these two timepoints, its development over the session should increase more under vagus than under sham stimulation.

### ERPs

We expect a main effect of stimulus condition on the amplitude of the ERP components: more positive amplitudes on oddballs than standards (P3 effect) and on target words in sentences with syntactic violations than on control sentences (P600 effect), as well as more negative amplitudes on target words in semantic violation than control sentences (N400 effect).

Further, no effect of stimulation on the P3/P600 amplitude would be taken as evidence against the LC/NE-P3-P600 hypothesis. Conversely, the following effects would suggest that the respective component (P3, P600) is linked to LC/NE activity: 1) Main effect of stimulation on the ERP amplitude (larger amplitude under vagus than sham stimulation), 2) interaction effect between stimulus type (oddballs/syntactic violations vs standards/control sentences) and stimulation (vagus vs sham) in that amplitudes on oddballs/syntactic violations are larger under vagus than under sham, and/or 3) significant differences in group comparison tests regarding the interaction between stimulus condition and stimulation.

Since conversely, the N400 has not been proposed to be linked to LC/NE activity, its amplitude should not be affected by stimulation, meaning neither an interaction between stimulus condition (semantic violation vs control) and stimulation (vagus vs sham) nor a main effect of stimulation on amplitudes in the N400 time window.

## Results

### Subjective ratings on stimulation effects

In line with the previous literature (see Giraudier et al., 2025), subjective ratings on the 7-point scale suggested that participants experienced little to no side effects during the stimulation (*M* = 1.35, see Table 2 below). The Wilcoxon signed rank test (ratings were not normally distributed) revealed almost no differences between stimulation conditions (all *p*s > .332). Only stinging was experienced significantly stronger under vagus than under sham stimulation (*p* = .025).

**Table 2.** Mean subjective ratings (standard deviation) for the stimulation side effects in the sham and vagus condition on the scale ranging from 1 (not at all) to 7 (very much).

|  | Sham | Vagus |
| --- | --- | --- |
| Headache | 1.35 (0.66) | 1.35 (0.89) |
| Nausea | 1.1 (0.5) | 1.03 (0.16) |
| Dizziness | 1.33 (0.88) | 1.25 (0.63) |
| Neck pain | 1.23 (0.48) | 1.23 (0.58) |
| Neck contraction | 1.3 (0.56) | 1.4 (0.59) |
| Stinging sensation | 1.45 (0.9) | 1.83 (1.17) |
| Ear irritation | 1.15 (0.43) | 1.3 (1.02) |
| Concentration | 2.5 (1.17) | 2.4 (1.22) |
| Fluctuation of feelings | 1.2 (0.72) | 1.2 (0.61) |
| Unpleasant feelings | 1.23 (0.62) | 1.5 (0.91) |

### Norepinephrine markers

#### Salivary alpha amylase levels

sAA levels in each condition over all three timepoints are displayed in Figure 2A below. There was no significant main effect of stimulation on sAA levels (F(1,216) = [0.005], *p* = .944). There was also no main effect of timepoint on sAA levels (F(2,216) = [0. 565], *p* = .569), meaning that overall sAA did not systematically rise or fall across the session (see Fig. 2A). This suggests that neither the stimulation nor time-on-task significantly altered sAA. Crucially, there was also no significant interaction between timepoint and stimulation (F(2,216) = [0.050], *p* = .951, all Tukey HSD comparison *p*’s > .95). Similarly, there was no significant difference in our exploratory measure of sAA change values between stimulation conditions regarding post1 (*t*(72) = −0.185, *p* = .854) or post2 (*t*(72) = −0.549, *p* = .585). Together, this suggests that the stimulation did not have any effect on the NE marker sAA level.

**Figure 2.**
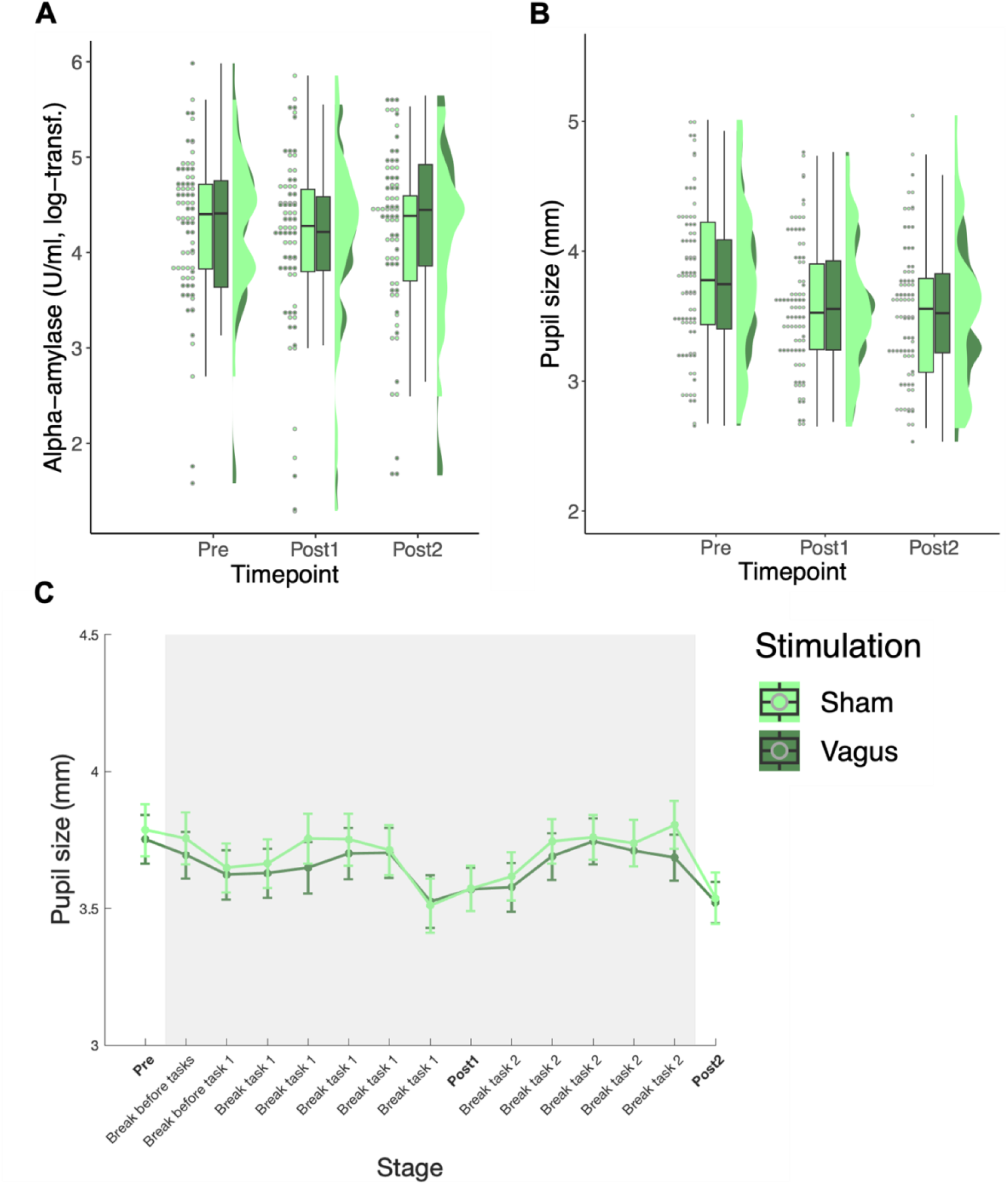
Results of NE markers. (**A**) sAA Levels and (**B**) baseline pupil size in each of the two stimulation conditions over the course of the three main timepoints: “pre” = baseline measurement before onset of stimulation, “Post1” = during stimulation, in break between tasks. “Post2” = right after tasks were finished and stimulation was turned off again. Raincloud plots depict individual data points (dots), their distribution (half–violin density), and summary statistics (boxplots). (**C**) depicts mean pupil size over the entire course of the experiment in each of the stimulation conditions. The grey box signifies the time window in which the stimulation was on and error bars indicate the standard error of the mean.

#### Baseline pupil size

Baseline pupil size in each stimulation condition over all three main timepoints is displayed in Figure 2B below. There was a significant main effect of timepoint ((F(2,234) = [4.476], *p* = .944) in that pupil size decreased from baseline to post2 (*p* = .015), indicating general habituation over the experiment. However, the stimulation did not affect baseline pupil size, neither as a main effect (F(1,234) = [0.061], *p* = .806) nor in interaction with timepoint (F(2,234) = [0.016], *p* = .984). As preregistered, we also explored potential more fine-grained effects of stimulation on pupil size. Figure 2C displays pupil size in all 15 measurements across the course of the entire experiment. However, we failed to find any shorter-lived effects, as paired t-tests revealed no significant differences between vagus and sham condition at any timepoint (all Bonferroni-corrected *p*’s > .551). Thus, the stimulation did not have any measurable effect on the NE marker baseline pupil size.

### Oddball task

#### Behavioral performance

Overall accuracy was very high (*M* = 96%; *SD* = 2%) and was significantly better in the standard condition (*M* = 99%, *SD* = 2%) than in the oddball condition (*M* = 84%, *SD* = 8%; *β* = −2.867, *SE* = 0.164, *z* = −17.471, *χ2* = 91.34, *p* < .001). The stimulation did not affect accuracy, neither as a main effect (*β* = 0.115, *SE* = 0.124, *z* = 0.928, *χ2* = 0.84, *p* = .359), nor in interaction with stimulus condition (*β* = −0.230, *SE* = 0.2, *z* = −1.150, *χ2* = 1.249, *p* = .263). Within correct trials, responses to standard stimuli were significantly faster than to oddball stimuli (*β* = 0.155, *SE* = 0.009, *t* = 17.306, *χ2* = 86.578, *p* < .001). However, there was no significant main effect of the stimulation on RTs (*β* = −0.002, *SE* = 0.011, *t* = −0.163, *χ2* = 0.027, *p* = .869). Lastly, there was no significant interaction between stimulus condition and stimulation on RTs in the oddball task (*β* = −0.001, *SE* = 0.011, *t* = −0.117, *χ2* = 0.014, *p* = .904). Note that all reported analyses regarding behavior were exploratory.

#### Effect of stimulus condition and stimulation on P3 amplitude

Stimulus condition means of the ERP amplitude in the oddball task are shown in Figure 3. We find the classic P3 effect: ERP amplitudes (300-600 ms) over posterior regions were significantly larger for oddballs (*M* = 9.96 mV, *SD* = 8.37) than standards (*M* = 3.93 mV, *SD* = 7.22; *β* = 6.041, *SE* = 0.512, *t* = 11.81, *χ2* = 60.829, *p* < .001). However, the stimulation did not significantly affect ERP amplitudes, neither as a main effect (*β* = −0.36, *SE* = 0.26, *t* = −1.382, *χ2* = 1.913, *p* = .167; mean condition difference = 0.33 mV) nor in interaction with stimulus condition (*β* = −0.002, *SE* = 0.27, *t* = −0.008, *χ2* = 5.877, *p* = .994). None of the preregistered posthoc group contrasts (stimulus condition x stimulation) revealed any significant effects (all *p*’s > .531). The additional, exploratory cluster-based permutation test also revealed no significant ERP amplitude differences between the vagus and sham condition (max cluster mass = 53.18; *p* = .212). Thus, taVNS did not measurably influence the P3.

**Figure 3.**
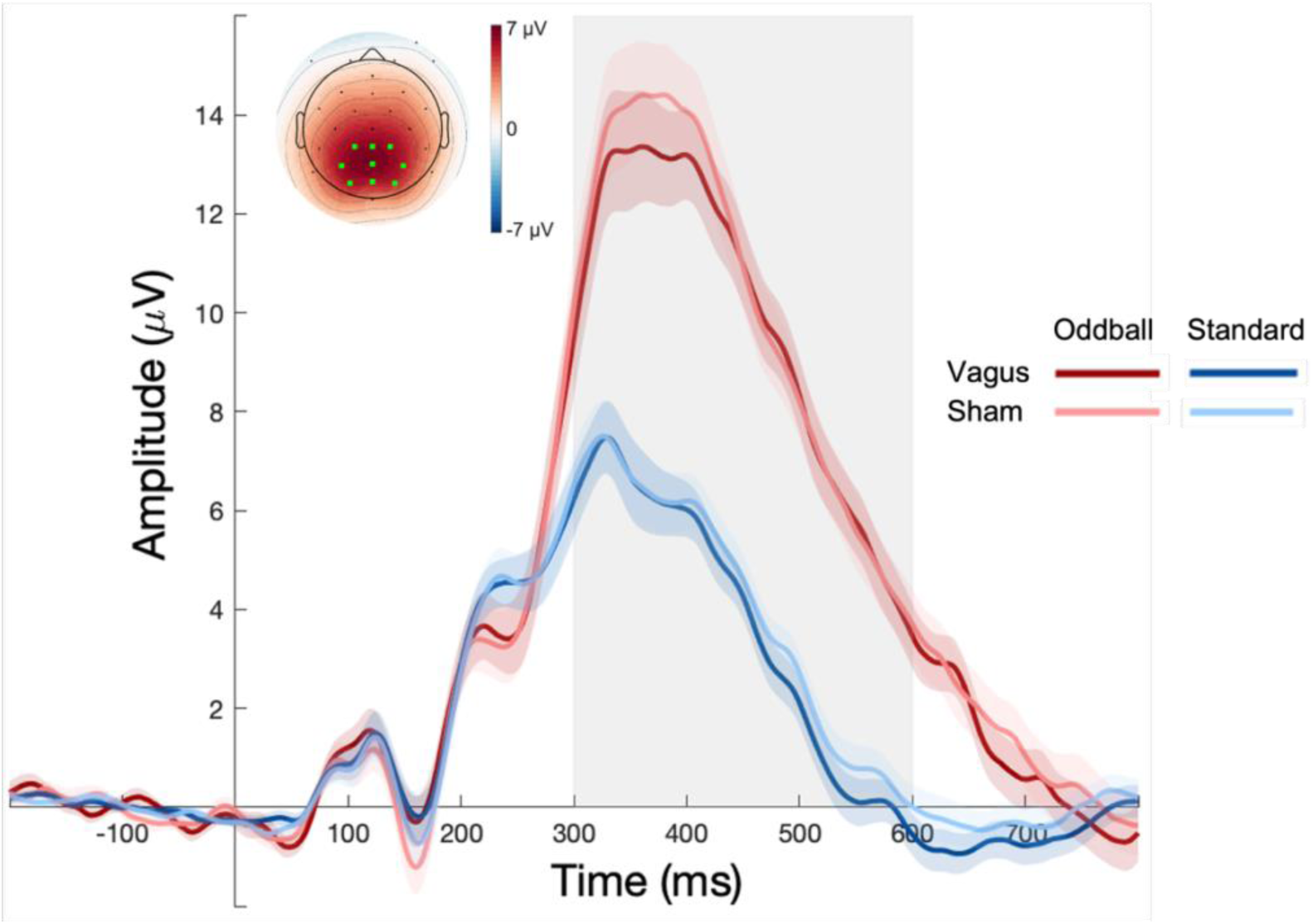
Grand averaged ERP amplitude for the four conditions in the oddball task. P3 amplitudes were calculated as the mean in the time interval marked by the grey box (300-600 ms) within the posterior channel cluster marked in light green in the topographic map. The topographic map shows the mean amplitude difference between oddball and standard stimuli within the same time window. Error bands indicate the standard error of the mean (SEM).

#### Influence of sAA change on stimulation effect on P3

Although the stimulation did not significantly affect sAA levels on a group level, we explored whether a potential effect of the stimulation on the P3 amplitude might be restricted to participants for which the stimulation increased sAA levels (as an index of LC/NE engagement). We quantified this by using sAA change values (see Method). The rationale was the following: If the stimulation in a session would successfully affect the LC/NE system, NE markers such as sAA levels should increase. We thus explored whether an (individual) increase of sAA level, possibly due to the stimulation, affected P3 amplitude. We did so by including the continuous measure of sAA change (two values per participant, one for sham and one for stimulation) as a predictor in the original P3 model, that is, in addition to an interaction with stimulus condition and stimulation. However, sAA change was not related to any effect of the stimulation on the P3. Regarding the change to post1 (during the stimulation), there was no significant main effect of sAA change on P3 amplitude (*β* = −0.232, *SE* = 0.294, *t* = −0.789, *χ2* = 0.526, *p* = .468) and no interaction between sAA and stimulus condition (*β* = 0.277, *SE* = 0.242, *t* = 1.140, *χ2* = 1.258, *p* = .262), or sAA and stimulation (*β* = −0.152, *SE* = 0.555, *t* = −0.273, *χ2* = 3.351, *p* = .341). Lastly, there was no 3-way interaction between sAA change, stimulus condition, and stimulation (*β* = −0.095, *SE* = 0.571, *t* = −0.167, *χ2* = 0.026, *p* = .871). The same pattern was found regarding the change to post2 (after the stimulation): There was no significant main effect of sAA change on P3 amplitude (*β* = 0.218, *SE* = 0.28, *t* = 0.778, *χ2* = 0.579, *p* = .447) and no significant interaction of sAA change with stimulus condition (*β* = 0.219, *SE* = 0.282, *t* = 0.777, *χ2* = 0.585, *p* = .445) or stimulation (*β* = 0.306, *SE* = 0.523, *t* = 0.586, *χ2* = 0.318, *p* = .573). Again, there was no 3-way interaction of sAA change with stimulus condition and stimulation (*β* = 1.076, *SE* = 0.542, *t* = 1.986, *χ2* = 3.823, *p* = .051). In sum, changes in sAA levels over the course of the experiment were unrelated to P3 amplitude, showing no main effect and no interaction with stimulation or stimulus condition.

#### Influence of task order on stimulation effect on P3

Lastly, we explored whether the stimulation effect on the P3 might have been dependent on stimulation duration (Bömmer et al., 2024; Giraudier et al., 2024). We thus investigated whether task order (i.e., whether the oddball task came earlier in the session or not) affected the stimulation effect on the P3 by including it as an additional predictor in the model. However, task order did not influence how the stimulation affected ERP amplitudes as there was no significant 3-way interaction between task order, stimulus condition, and stimulation (*β* = −0.906, *SE* = 0.541, *t* = −1.676, *χ2* = 2.81, *p* = .1) and no 2-way interaction between task order and stimulation (*β* = −0.670, *SE* = 0.516, *t* = −1.298, *χ2* = 1.735, *p* = .188).

### Sentence processing task

#### Behavioral performance

Overall accuracy in the sentence acceptability judgement was high (*M* = 93%; *SD* = 4%), indicating that participants attentively read the sentences. Accuracy was significantly better regarding correct control sentences (*M* = 94%, *SD* = 4%) than sentences including violations (*M* = 92%, *SD* = 5%; *β* = −0.362, *SE* = 0.133, *z* = −2.730, *χ2* = 6.81, *p* = .009).

However, the stimulation did not affect accuracy, neither as a main effect (*β* = 0.138, *SE* = 0.103, *z* = 1.337, *χ2* = 1.688, *p* = .194), nor in interaction with stimulus condition (*β* = −0.124, *SE* = 0.152, *z* = −0.815, *χ2* = 0.626, *p* = .429). Within correct trials, response times were neither affected by stimulus condition (*β* = −0.116, *SE* = 0.060, *t* = −1.92, *χ2* = 3.614, *p* = .057), nor stimulation (*β* = 0.028, *SE* = 0.062, *t* = 0.445, *χ2* = 0.203, *p* = .653), nor their interaction (*β* = −0.030, *SE* = 0.045, *t* = −0.678, *χ2* = 0.461, *p* = .497). Note that all reported analyses regarding behavior were exploratory and collapsed across violation types.

#### Effect of stimulus condition and stimulation on P600 amplitude

Mean ERP amplitudes over time in each of the stimulus conditions in the sentence processing task are shown in Figure 4. Overall, we found a syntactic as well as semantic^2^ P600 effect. Amplitudes between 600-900 ms at parietal sites were significantly larger in response to syntactic violations (*M* = 4.91 mV, *SD* = 8.48) than respective correct controls (*M* = 0.677 mV, *SD* = 7.55; *β* = 4.198, *SE* = 0.479, *t* = 8.755, *χ2* = 43.42, *p* < .001). Analogously, P600 amplitudes were significantly larger on semantic violations (*M* = 2.81 mV, *SD* = 7.89) than respective correct controls (*M* = 1.8 mV, *SD* = 7.16; *β* = 0.977, *SE* = 0.23, *t* = 4.252, *χ2* = 15.196, *p* < .001).

**Figure 4.**
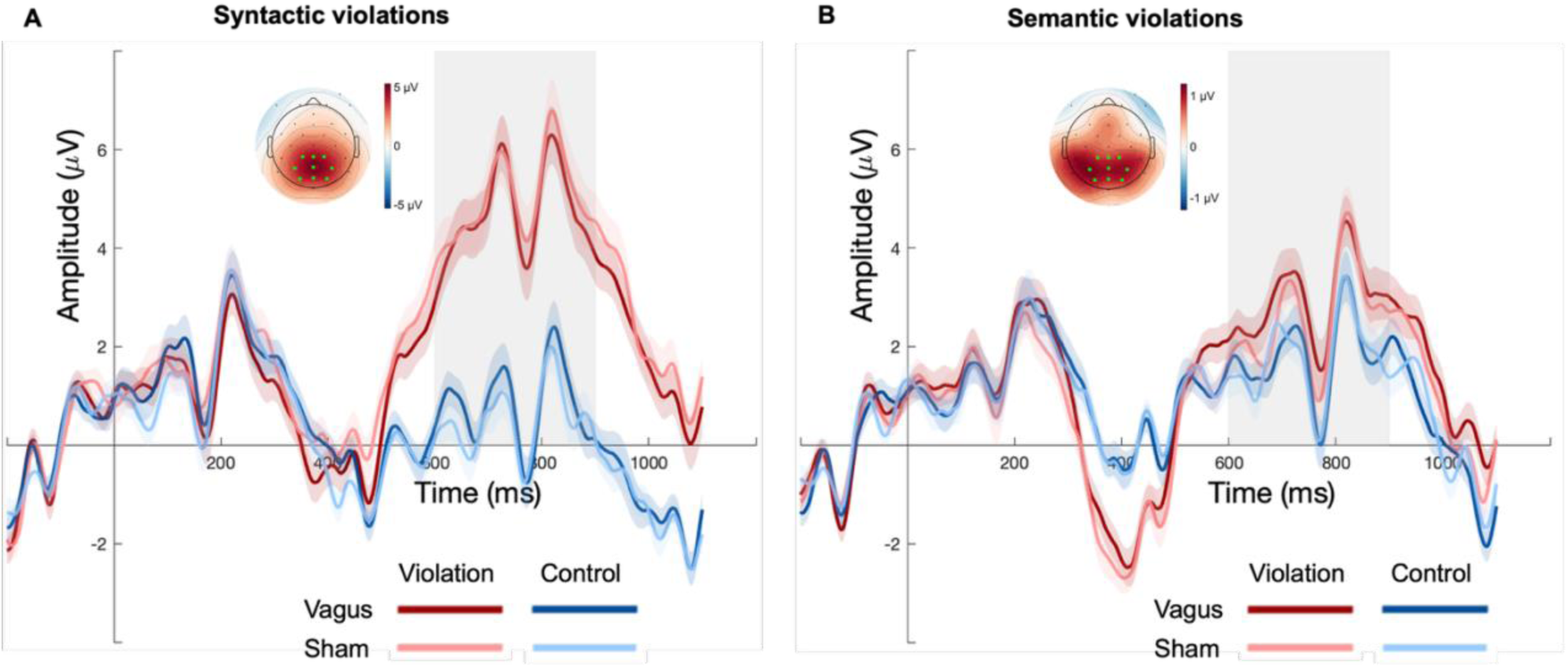
Grand averaged ERP amplitudes in the sentence processing task aligned to target word onset for the (**A**) syntactic and (**B**) semantic sentence type, each divided by stimulation condition. P600 amplitudes were calculated as the mean in the time interval marked by the grey box (600-900 ms) within the posterior channel cluster marked in light green in the topographic maps. Topographic maps show the mean amplitude difference between violation and control stimuli within the same time window. Error bands indicate the standard error of the mean.

Regarding the syntactic P600, the stimulation did not significantly affect ERP amplitudes, neither as a main effect (*β* = 0.013, *SE* = 0.244, *t* = 0.051, *χ2* = 0.003, *p* = .959, mean condition difference = 0.01 mV) nor in interaction with stimulus condition (*β* = −0.586, *SE* = 0.395, *t* = −1.484, *χ2* = 2.201, *p* = .138). None of the preregistered posthoc group contrasts (condition x stimulation) revealed any significant effects (all *p*’s > .760). Likewise, the additional, exploratory cluster-based permutation test also did not reveal any significant ERP amplitude differences between the vagus and sham condition (max cluster mass = 20.88; *p* = .67).

The same results pattern was obtained regarding the semantic P600: There was no significant main effect of stimulation (*β* = 0.192, *SE* = 0.190, *t* = 1.007, *χ2* = 1.015, *p* = .314, mean condition difference = 0.18 mV), no interaction between stimulus condition and stimulation (*β* = 0.275, *SE* = 0.381, *t* = 0.723, *χ2* = 0.522, *p* = .47), and no significant posthoc group contrasts (all *p*’s > .659). Again, the additional, exploratory cluster-based permutation test did not reveal any significant ERP amplitude differences between the vagus and sham condition (max cluster mass = 20.19; *p* = .692). Thus, taVNS did not measurably influence the amplitude of the syntactic or semantic P600.

#### Influence of sAA change on stimulation effect on P600

As in the oddball task, we explored whether a potential effect of the stimulation on the P600 amplitude might be restricted to participants for which the stimulation increased sAA levels (indicating LC/NE engagement). To that end, we included the continuous measure of sAA change as an interacting predictor in the original P600 models in addition to stimulus condition and stimulation.

Regarding the syntactic violations, sAA change to post1 (during the stimulation) did not significantly affect P600 amplitudes (*β* = 0.055, *SE* = 0.235, *t* = 0.234, *χ2* = 0.106, *p* = .744). There were also no significant interactions between sAA change and stimulation (*β* = - 0.786, *SE* = 0.481, *t* = −1.637, *χ2* = 2.745, *p* = .098), or stimulus condition (*β* = −0.406, *SE* = 0.337, *t* = −1.203, *χ2* = 1.42, *p* = .233). Lastly, there was no significant 3-way interaction between sAA change, stimulus condition, and stimulation (*β* = −0.253, *SE* = 0.775, *t* = −0.326, *χ2* = 0.116, *p* = .734). For the change to post2 (after the stimulation), again, there was no significant main effect of sAA change (*β* = 0.362, *SE* = 0.205, *t* = 1.770, *χ2* = 3.127, *p* = .077) or interaction between sAA change and stimulation (*β* = −0.397, *SE* = 0.457, *t* = −0.869, *χ2* = 0.71, *p* = .4). Notably though, sAA change to post2 did significantly interact with stimulus condition (*β* = −1.012, *SE* = 0.390, *t* = −2.594, *χ2* = 6.665, *p* = .01), reflecting the fact that P600 amplitudes were larger the more sAA levels increased, but only in response to violations (*β* = 1.017, *SE* = 0.321, *p* = .002), not controls (*β* = 0.005, *SE* = 0.293, *p* = .987). Lastly, there was no significant 3-way interaction between sAA change, stimulation, and stimulus condition (*β* = −0.369, *SE* = 0.756, *t* = −0.488, *χ2* = 0.248, *p* = .051)

Regarding semantic violations and post1 (during the stimulation) there was no significant main effect of sAA change (*β* = −0.265, *SE* = 0.194, *t* = −1.368, *χ2* = 1.945, *p* = .163), no interaction of sAA and stimulation (*β* = −0.105, *SE* = 0.396, *t* = −0.265, *χ2* = 0.073, *p* = .787) and no interaction of sAA change and stimulus condition (*β* = 0.202, *SE* = 0.277, *t* = 0.727, *χ2* = 0.548, *p* = .459) on P600 amplitude. There was also no significant 3-way interaction of sAA change with stimulation and stimulus condition (*β* = 0.058, *SE* = 0.610, *t* = 0.095, *χ2* = 0.008, *p* = .926). In the same vein, sAA change to post2 (after the stimulation) did not significantly affect P600 amplitude (*β* = −0.027, *SE* = 0.261, *t* = −0.104, *χ2* = 0.009, *p* = .923) and again did not significantly interact with stimulation (*β* = 0.110, *SE* = 0.378, *t* = 0.292, *χ2* = 0.076, *p* = .783), or with stimulus condition (*β* = 0.236, *SE* = 0.311, *t* = 0.759, *χ2* = 0.534, *p* = .465). Lastly, there was also no significant 3-way interaction between sAA change, stimulus condition, and stimulation (*β* = 0.751, *SE* = 0.612, *t* = 1.228, *χ2* = 1.492, *p* = .222).

Thus, sAA change did not influence any effect of the stimulation on the syntactic or semantic P600. However, the observed exploratory interaction between sAA change and P600 amplitude indicates a relationship between the syntactic P600 and a marker of NE independently of the stimulation.

#### Influence of task order on stimulation effect on P600

We again explored the possibility of time-dependent effects of stimulation. We thus included task order as an additional (interacting) predictor in the P600 models. However, task order did not influence how the stimulation affected P600 amplitudes as there was no significant 3-way interaction between task order, stimulus condition, and stimulation (syntactic P600: *β* = 0.147, *SE* = 0.78, *t* = 0.186, *χ2* = 5.358, *p* = .253; semantic P600: *β* = 1.136, *SE* = 0.760, *t* = 1.494, *χ2* = 2.226, *p* = .136). Likewise, there was no 2-way interaction between task order and stimulation (syntactic P600: *β* = −0.261, *SE* = 0.493, *t* = −0.529, *χ2* = 0.294, *p* = .588; semantic P600: *β* = −0.072, *SE* = 0.380, *t* = −0.189, *χ2* = 0.035, *p* = .851).

#### Effect of condition and stimulation on N400 amplitude

As a control component that is not presumed to be linked to NE release, we preregistered to test whether the N400 is affected by the stimulation. At the parietal channel cluster (same ROI as P3 and P600, see methods) we replicated the classic N400 effect: Amplitudes between 300-500 ms were significantly more negative on semantic violations (*M* = −0.061 mV, *SD* = 7.45) than respective correct controls (*M* = 0.612 mV, *SD* = 7.32; *β* = − 0.680, *SE* = 0.214, *t* = −3.182, *χ2* = 9.241, *p* = .002, see Fig. 4B). As expected, the stimulation did not significantly affect N400 amplitudes, neither as a main effect (*β* = 0.132, *SE* = 0.18, *t* = 0.695, *χ2* = 0.483, *p* = .487, mean condition difference: 0.136 mV) nor in interaction with stimulus condition (*β* = 0.398, *SE* = 0.38, *t* = 1.047, *χ2* = 1.098, *p* = .295). None of the preregistered posthoc group contrasts (stimulus condition x stimulation) revealed any significant effects (all *p*’s > .601).

We found the identical pattern of results at the centro-parietal channel cluster (common N400 ROI, see methods): There was an N400 effect between 300-500 ms in that amplitudes were significantly more negative on semantic violations (*M* = −0.268 mV, *SD* = 8.44) than correct controls (*M* = 0.517 mV, *SD* = 8.34; *β* = −0.8, *SE* = 0.246, *t* = −3.255, *χ2* = 9.631, *p* = .002). Again, the stimulation did not significantly affect N400 amplitudes, neither as a main effect (*β* = 0.3, *SE* = 0.216, *t* = 0.695, *χ2* = 1.385, *p* = .174, mean condition difference: 0.299 mV) nor in interaction with stimulus condition (*β* = 0.602, *SE* = 0.433, *t* = 1.390, *χ2* = 1.935, *p* = .164). None of the preregistered posthoc group contrasts (stimulus condition x stimulation) revealed any significant effects (all *p*’s > .206).

### Relationship between ERP components across tasks

Lastly, we explored a potential relationship between the P600 and P3 effect (i.e., the mean amplitude difference between experimental and control condition) within participants, which would be predicted if both relied on the same neural generator.

Indeed, the syntactic P600 effect was positively predicted by the P3 effect (*β* = 0.412, *SE* = 0.141, *t* = 2.916, *p* = .005, Figure 5). This relationship was independent of the stimulation, as there was no interaction with stimulation (*β* = 0.105, *SE* = 0.202, *t* = 0.518, *p* = .606). Further, the relationship was unlikely due to more general cognitive state or mood, as the correlation persisted across both sessions (i.e., P600 effect in session 1 correlated with P3 effect in session 2 and vice versa, both *p*’s < .002). Thus, we found that participants with larger P3 effects in the oddball task also displayed larger syntactic P600 effects in the sentence processing task independent of the stimulation, session, and order of tasks, further linking the two late, positive components.

**Figure 5.**
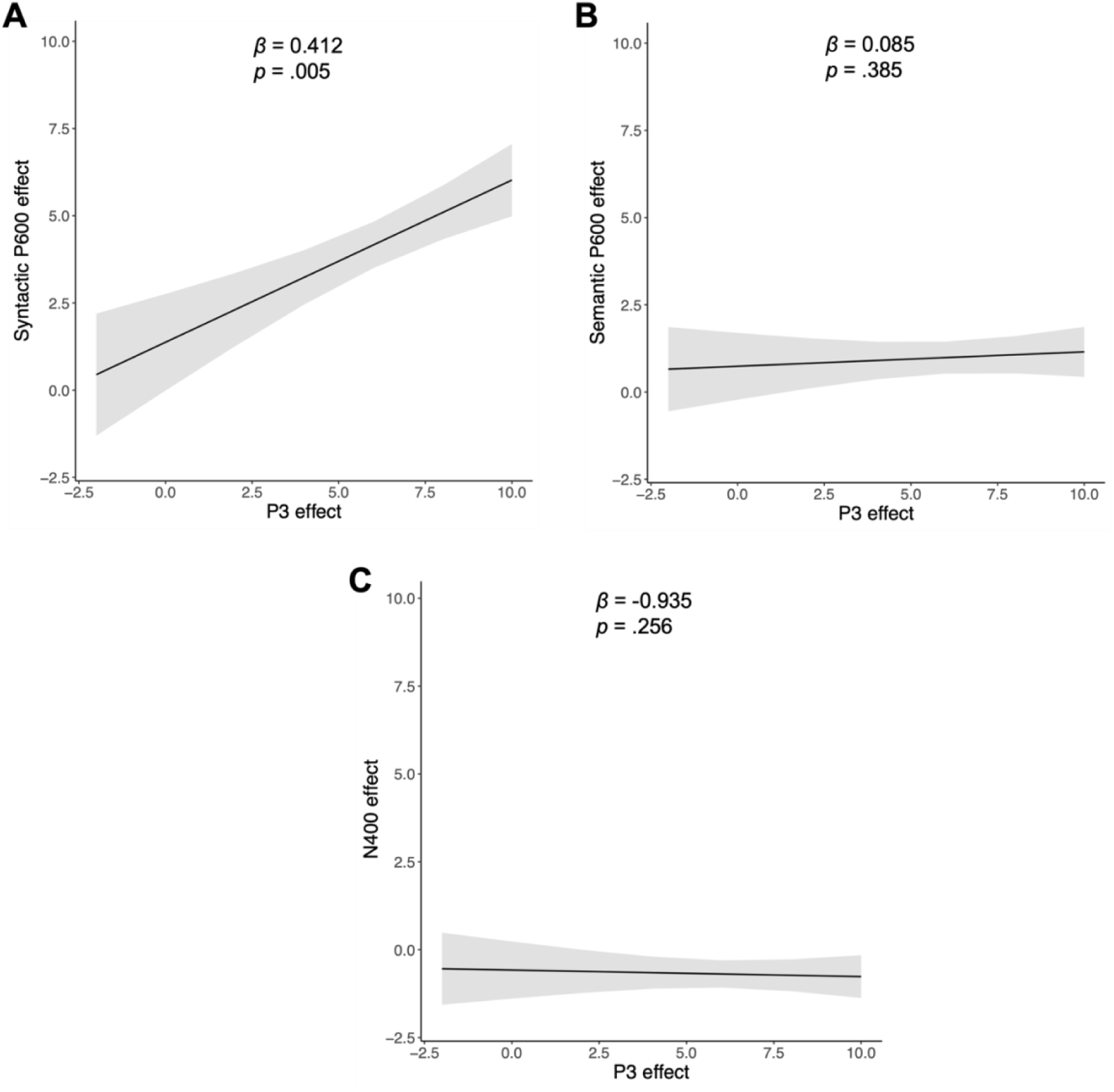
Relationship (i.e., slope of fitted linear models) between P3 effect and (**A**) syntactic P600 effect, (**B**) semantic P600 effect, and (**C**) N400 effect in response to semantic violations. Shadows represent 95% confidence intervals.

Notably, this relationship of the P3 and P600 effect was specific to the syntactic P600 effect, as we did not find such an effect for the semantic P600 effect (main effect: (*β* = 0.085, *SE* = 0.098, *t* = 0.873, *p* = .385; interaction with stimulation: *β* = −0.088, *SE* = 0.14, *t* = −0.630, *p* = .530, Figure 5 below) or the N400 effect in response to semantic violations (main effect: *β* = −0.935, *SE* = 0.817, *t* = −1.144, *p* = .256; interaction with stimulation: *β* = 0.087, *SE* = 0.119, *t* = 0.734, *p* = .465, Figure 5).

## Discussion

The present study investigated a modulatory effect of continuous transcutaneous auricular vagus stimulation on the ERP components P3 and P600, both of which have been proposed to reflect phasic NE activity in the locus coeruleus (e.g., Nieuwenhuis et al., 2005; Sassenhagen & Bornkessel-Schlesewsky, 2015). Moving beyond correlational evidence (e.g., Contier et al., 2024), we set out to test the hypothesis that phasic NE release causally contributes to the generation of the two late, positive ERP components. Using a within-subject, between-session design, we applied continuous taVNS to the cymba conchae, which has been linked to activation of the LC/NE system, and compared it to sham stimulation at the earlobe. During stimulation in both sessions, participants completed an active visual oddball task (to elicit the P3) and an active sentence processing task including both syntactic and semantic violations (to elicit the P600). We observed robust main effects of stimulus condition: Oddball stimuli elicited larger P3 amplitudes than standard stimuli (P3 effect), and both syntactic and semantic violations elicited larger P600 amplitudes than target words in corresponding control sentences (P600 effect). In contrast, taVNS had no measurable effect on the amplitude of either component, neither by itself nor in interaction with stimulus condition. Notably, these null effects must be understood in the context of a possibly ineffective neuromodulatory manipulation. To assess whether taVNS affected NE as intended, we additionally measured two well-established physiological markers of NE activity (e.g., Joshi et al., 2016; Laeng et al., 2012; Nater & Rohleder, 2009): salivary alpha-amylase (sAA) and baseline pupil size at three timepoints (before, during, and after the stimulation). However, neither of the two measures showed evidence of change specifically due to the stimulation in the current study. Thus, our manipulation checks do not allow us to draw any major conclusions about a potential causal link between the two ERP components and NE.

The absence of taVNS effects on the ERP components may also relate to the mode of stimulation. The proposed link between NE and both the P3 and P600 rests on the assumption that these components primarily reflect phasic, rather than tonic, LC/NE activity. In contrast, the present study employed continuous taVNS, which may preferentially influence tonic LC/NE levels or lead to habituation over time. This choice was guided by empirical and methodological considerations. First, prior taVNS studies using continuous stimulation reported modulatory effects on the P3 component (e.g., Ventura-Bort et al., 2018, 2021; Giraudier et al., 2024). Second, at the time of study design, intermittent protocols (e.g., 30 s on / 30 s off) represented the most common alternative (e.g., Lewine et al., 2019; C. V. Warren et al., 2020; *tVNS Technologies GmbH* has also embedded this on/off cycle in their commercial device), but these are not truly phasic either, as stimulation is not time-locked to individual task events and LC responses to VNS appear tightly coupled to stimulation epochs. While brief, event-locked stimulation bursts may provide a more direct means of targeting phasic LC/NE responses, such protocols also pose challenges for ERP research, as stimulation-related somatosensory and electrical artefacts would be temporally aligned with the ERP components of interest. Finally, from a theoretical perspective, continuous stimulation could modulate phasic responsiveness in different directions depending on participants’ baseline arousal. According to the inverted U-shaped relationship between tonic and phasic LC/NE activity (Aston-Jones & Cohen, 2005), moderate tonic activation may facilitate phasic responses, whereas higher tonic levels may suppress them. Continuous taVNS may therefore have shifted participants differentially along this function, which potentially obscured consistent group-level effects on ERP components. This variability in participants’ underlying arousal state could account for the heterogeneous effects on the P3 reported with continuous taVNS (e.g., d’Agostini et al., 2022; Fischer et al., 2018; Giraudier et al., 2024; Ventura-Bort et al., 2018). In future work, stimulation could be adjusted individually by incorporating simple physiological measures such as baseline pupil size or brief test bursts that elicit small, measurable pupil responses, helping to place participants at comparable arousal levels. Such physiologically guided, individualized calibration may help reduce between-participant differences in tonic NE state and improve the consistency of neuromodulatory engagement. Whether continuous taVNS impacts phasic ERP components may further depend on the experimental context (e.g., properties of stimuli, stimulus context, task-relevance). For instance, continuous stimulation might exert stronger effects on late positive ERPs when phasic responses are elicited by emotionally salient (i.e., arousing) stimuli (e.g., Ventura-Bort et al., 2025) or target stimuli are presented in the context of arousing stimuli (e.g., Ventura-Bort et al., 2018; Giraudier et al., 2024; for discussion on the sAA, see Giraudier et al., 2022), or when only target stimuli require a behavioral response (c.f., Ventura-Bort et al., 2018, Giraudier et al., 2024; in contrast to the present study, in which a button press was required for both target and standard stimuli). Thus, the modulation of these ERP components by taVNS may be shaped by the interaction between stimulation mode and the arousing properties of the stimuli, or context (which also likely includes the saliency of stimuli given by task relevance).

The present findings thus highlight crucial methodological considerations but also the importance of the stimulation pattern, which may include settings to target phasic responses specifically. Recent advances in short-term taVNS protocols that deliver brief bursts of stimulation have been proposed as a more precise and temporally targeted method for specifically affecting the phasic LC/NE response and have repeatedly been shown to enhance pupil dilation (e.g., d’Agostini et al. 2023; Lloyd et al., 2023; Ludwig et al., 2023; Pervaz et al., 2025; Sharon et al., 2021; Skora et al., 2024; Villani et al., 2022). Importantly, this can be used time-locked to cognitive events, specifically targeting the processing of task-relevant stimuli. For example, short bursts of taVNS during an encoding task led to better memory and larger pupil dilation for negative events (Ludwig et al., 2025). This method has also been shown to enhance target detection in rodents (Mridha et al., 2021). Interestingly, Mridha and colleagues (2021) observed the strongest phasic, stimulation evoked pupillary and neural responses during intermediate tonic NE activity, supporting the notion that providing baseline arousal conditions is crucial to successfully manipulate phasic NE activity. This phasic, event-locked method may recruit the LC/NE system more reliably than continuous, tonic stimulation approaches, offering promising methods to further investigate NE contributions to ERP components such as the P3 and P600. Such bursts could be delivered time-locked to each stimulus, for example shortly before every trial in the oddball task or before the critical word in each sentence. This would differ from continuous stimulation by restricting neuromodulatory input to discrete, task-relevant moments. However, several practical challenges may also come with this approach. First, the optimal timing of a burst relative to the cognitive processes generating the P3 and P600 and behavioral response is non-trivial.

Second, both tasks – especially the oddball task – present stimuli at relatively high frequency, meaning that trial-wise bursts would form a dense stimulation schedule that might approximate continuous stimulation, with possible habituation effects also compromising the direct comparison of both paradigms. Third, brief bursts delivered around stimulus onset may introduce stimulation condition-related somatosensory and/or electrical artefacts that are time-locked to the ERP components of interest, complicating interpretation of any amplitude changes. Nevertheless, event-locked protocols remain promising for future work aiming to selectively modulate phasic LC/NE activity. In this context, it would also be highly informative to include task-evoked (phasic) pupil data in addition to the baseline (tonic) pupil measures as reported here. Phasic pupillary responses are closely linked to transient LC/NE activation (e.g., Joshi & Gold, 2016) and would therefore provide a complementary index of whether taVNS specifically modulated phasic NE activity. Future studies combining event-locked taVNS with continuous pupillometry may thus help clarify the precise relationship between stimulation-induced LC/NE engagement and ERP components such as the P3 and P600. Ultimately, these considerations underscore the need for standardized, non-invasive stimulation and recording protocols to target phasic NE activity specifically.

Given the likely complex relationship between positive ERP components and the neuromodulatory brain stem system, taVNS studies should furthermore be complemented by other modes of intervention. One avenue for future research might be to use alternative methods to stimulate involved peripheral nerves such as the greater occipital nerve and the trigeminal nerve (e.g., Luckey et al., 2023; van Boekholdt et al., 2021) as well as pharmacological manipulations using NE agonists that have already been shown to modulate the P3 (e.g., Joseph & Sitaram, 1989). Another promising method may be isometric handgrip, a form of static muscular contraction in which participants constantly squeeze a handgrip device over several seconds (e.g., Nielsen & Mather, 2015). This transiently elevates sympathetic arousal and has been shown to increase phasic LC responsivity as well as pupil dilation in response to salient events and improve subsequent cognitive performance (Mather et al., 2020). As a brief, portable, and non-invasive technique, the handgrip method may serve as a valuable complementary tool here.

Although our taVNS protocol did not modulate ERP amplitudes or physiological NE markers, we did find two notable pieces of exploratory evidence supporting the P600’s link to both the P3 and NE, respectively. First, we observed a within-subject correlation between the P3 effect and syntactic P600 effect that was independent of taVNS stimulation and session. Together with previous evidence on their shared link to the pupillary response (Contier et al., 2024), reaction time (Sassenhagen & Bornkessel-Schlesewsky, 2015) and the similarity of their waveforms (Sassenhagen & Fiebach, 2019), this correlation of ERP effects supports the possibility of shared or overlapping neurocognitive mechanisms, such as salience detection or context updating, potentially modulated by neuromodulators like NE. Secondly, larger sAA increase over the course of the experiment was associated with larger syntactic P600 amplitudes, suggesting that endogenous NE activity, as indexed by sAA, may enhance the neural processes underlying cognitive processing in linguistic contexts. Importantly, this exploratorily analysed relationship was selective to violation trials, suggesting that LC/NE activity may specifically support processing in situations of conflict and/or uncertainty. Although correlational, this result supports the notion that the P600 is sensitive to NE and suggests that naturally occurring variation in NE activity may be an important factor in late, positive ERP amplitude variation, even in the absence of experimental modulation. Lastly, these two links (i.e., the link of the P600 to the P3 and to the NE marker sAA level) were both found specifically for the P600 in response to syntactic, but not semantic violations. This may be partly attributable to the comparatively smaller semantic P600 effect (compared to the syntactic P600 effect), which likely constrained amplitude variability necessary to reveal reliable correlations with other neural or physiological measures. Nevertheless, it is an open question whether the P600s in response to syntactic and semantic violations reflect the same process and whether both or only the former are related to the P3 (Leckey & Federmeier, 2019). Thus, both findings certainly stimulate more research into the possible differences between the semantic and syntactic P600 and encourage incorporating a more diverse set of linguistic anomalies into single P600 studies to potentially uncover unknown processing differences and potential differential relationships to the P3 and NE.

The present pattern of the P3’s and P600’s relationship to sAA levels is also consistent with models in which both the P3 and P600 reflect phasic LC/NE responses whose magnitude depends nonlinearly on the tonic NE state. Because tonic NE modulates phasic responsiveness in an inverted-U-shaped fashion (see above), tonic ERP relationships may be positive, negative, or absent across individuals. This variability is well documented for the P3 (e.g., Chung et al., 2024; Contier et al., 2024; Murphy et al., 2011), and our finding that the syntactic P600 covaries with sAA similarly suggests that its phasic expression is likewise shaped by tonic NE mode. Thus, mixed associations with tonic markers are expected, yet the presence of such relationships in at least some cases supports the possibility that late positive ERP components are influenced by LC/NE activity.

In conclusion, our study finds no influence of continuous taVNS on either the P3 or P600 ERP component nor on physiological markers of NE, suggesting that the stimulation protocol may not have effectively modulated the LC/NE system. Thus, our results cannot provide evidence for a causal role of NE in the neural generation of the P3 or P600, but they also do not falsify this hypothesis. Instead, they highlight the need for more standardized taVNS protocols. Importantly, they emphasize the need for better-targeted manipulations of phasic NE activity, such as phasic, stimulus-aligned taVNS to advance our understanding of potential links between the late, positive ERP components and the LC/NE system. Lastly, a link between the P600 and P3 as well as increased sAA observed in exploratory analyses supports the view that the two positivities may be related and phasic NE release may contribute to late-stage language processing, particularly in response to incongruencies.

## Acknowledgements

We thank L. Y. Lam, A. Sterzik, J. Kirschner, and J. Barthelmeß for their help with data collection. Funding was provided by the Research Focus Cognitive Sciences of the University of Potsdam and by the Deutsche Forschungsgemeinschaft (DFG, German Research Foundation, Project ID 317633480, SFB1287).

## Footnotes

1 Coding of relevant regressors: Stimulation: sham (-.5) vs. vagus (.5); Oddball task: standard (-.5) vs. oddball (.5); Sentence processing task: violation (-.5) vs. control (.5).

2 Note that our preregistration focused on the syntactic P600 and how it is potentially influenced by the stimulation. However, since we also observed a P600 regarding semantic violations, we also explored whether the semantic P600 is influenced by the stimulation.

## Supplementary material

**Figure A1.**
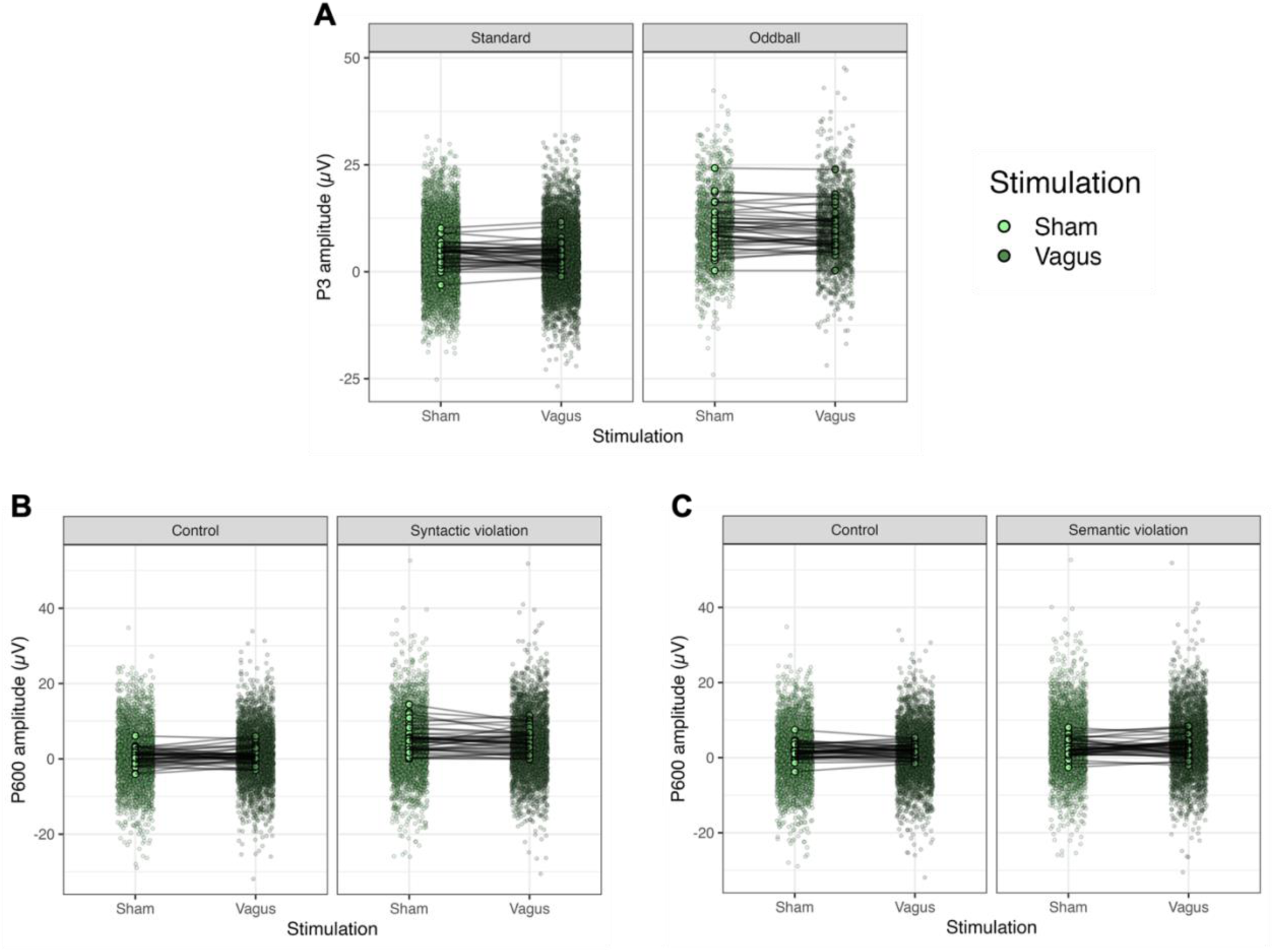
Trial-wise ERP amplitudes and participant-wise condition means for the (**A**) P3, (**B**) syntactic P600, and (**C**) semantic P600. Each point represents a single trial (jittered for visibility), and larger filled circles indicate the mean amplitude for each participant in each stimulation condition. Lines connect the means of the two stimulation conditions within participants. Panels are faceted by stimulus type (Standard vs. Oddball). Plots are provided for descriptive purposes and to illustrate the relative magnitude of inter-trial and inter-individual variability in the data entering the mixed-effects models.

